# Conformational Ensemble of Monomeric *α*-Synuclein in Aqueous and Crowded Environments as revealed by Markov State Model

**DOI:** 10.1101/2022.02.20.481191

**Authors:** Sneha Menon, Jagannath Mondal

## Abstract

140-residue intrinsically disordered protein *α*-synuclein (*α*S) is known to be susceptible to environmental cues/crowders and adopts conformations that are vastly variable in the extent of secondary structure and tertiary interactions. Depending upon the nature of these interactions, some of the conformations may be suitable for its physiological functions while some may be predisposed to aggregate with other partners into higher ordered species or to phase separate. However, the inherently heterogenous and dynamic nature of *α*S has precluded a clear demarcation of its monomeric precursor between aggregation-prone and functionally relevant aggregation-resistant states. Here, we optimally characterise a set of metastable conformations of *α*S by developing a comprehensive Markov state model (MSM) using cumulative 108 *µ*s-long all-atom MD simulation trajectories of monomeric *α*S. Notably, the dimension of the most populated metastable (85%) state (*R_g_* ∼ 2.59 (±0.45) nm) corroborates PRENMR studies of *α*S monomer and undergoes kinetic transition at 0.1-150 *µ*s time-scale with weakly populated (0.06%) random-coil like ensemble (*R_g_* ∼ 5.85 (±0.43) nm) and globular protein-like state (14%) (*R_g_* ∼ 1.95 (±0.08) nm). The inter-residue contact maps identify a set of mutually interconverting aggregation-prone *β*-sheet networks in the NAC region and aggregation-resistant long-range interactions between N- and C-terminus or helical conformations. The presence of crowding agents compacts the MSM-derived metastable conformations in a non-monotonic fashion and skews the ensemble by either introducing new tertiary contacts or reinforcing the innate contacts to adjust to the excluded-volume effects of such environments. These observations of crucial monomeric states would serve as important steps towards rationalising routes that trigger *α*S-associated pathologies.

**Significance statement:** *α*-synuclein, a neuronal protein, is often associated with neurogenerative diseases due to its tendency to self-assemble into higher ordered aggregates. While the monomeric precursor of this protein is intrinsically disordered, it is also known to be susceptible to biological environmental cues and adopts a wide range of conformations that are either primed for aggregation or remain in auto-inhibitory states. However, the inherently heterogenous nature of the monomeric form has prevented a clear dissection of aggregation-prone and functionally relevant aggregation-resistant states. Here, we resolve this via an atomistic characterisation of an optimal set of crucial metastable monomeric conformations via statistical modelling of computer simulated data. The investigation also sheds light on crowding-induced modulation of the ensemble and eventual fibrillation pathways.

## Introduction

*α*-Synuclein (*α*S) is a 140-residue, 14 kDa protein abundantly present in the brain, mainly in the presynaptic nerve terminals.^1^ The potential physiological roles attributed to *α*S include release of neurotransmittors and synaptic plasticity.^2, 3^ It is a prototypical intrinsically disordered protein (IDP) and its excessive accumulation is associated with the neurodegenerative Parkinson’s disease and other synucleinopathies.^4, 5^ Being natively unstructured, *α*S has the tendency to accumulate a variety of rapidly interconverting conformations under physiological conditions. ^6^ Some of these conformations are also susceptible to self-assemble into higher ordered species such as oligomers, protofibrils and fibrils across different length and time scales in a hierarchical process known as *amyloidogenesis*.^7, 8^ These ordered aggregates form intracellular inclusions known as Lewy bodies, which is a hallmark of these neurodegenerative diseases.^4^

Characterisation using a number of biophysical techniques has revealed that monomeric *α*S can either be natively unstructured in aqueous solution or alternatively adopt ordered structures under the influence of different conditions. Experimental studies indicate that endogenous cellular *α*-synuclein exists largely as *α*-helically folded tetramer that is resistant to aggregation.^9, 10^ It is postulated that destabilization of this physiological tetramer is a pre-requisite for *α*S misfolding and aggregation into abnormal oligomeric and fibrillar assemblies. In contrast, the aggregated states (oligomers and fibrils) are identified with extensive *β*-sheet content.^11, 12^ The thermokinetic models^13–15^ that describe amyloid formation predict a primary nucleation step in which aggregation-competent monomers of *α*S favorably coalesce into heterogenous oligomers of variable size and *β*-sheet content. These oligomers progressively evolve by monomer addition into protofilaments and thereby into mature amyloid fibrils. Different structure determination techniques have reported the structure of multiple polymorphs of *α*S fibril.^16–21^ EPR data and ssNMR studies have identified the amyloid core consisting of residues 30 to 100 forming parallel, in-register *β*-sheets stacked with other monomer units to form a fibril.

However, the cellular milieu is more complex than aqueous media and mostly involves a high concentration of *crowders*, whose impact has traditionally been realised in various properties of proteins such as folding, stability, ligand-binding, self-assembly.^22–25^ The crowded intracellular environment can have altering effects on protein aggregation and fibrillation, therefore it is now a standard practice to include crowding effects mimicking the cellular milieu in both experimental and theoretical studies. A number of earlier studies have reported that asyn fibrillation is accelerated in the presence of macromolecular crowding agents in vitro.^26–30^ Recent reports demonstrate the liquid-liquid phase separation (LLPS) of *α*S and postulate that this process is a critical event in the early lag phase preceding the aggregation process.^31, 32^ Intriguingly, the phase separation process has been found to be mediated by the presence of *molecular crowders* that increases the local protein concentration and facilitates condensation and droplet formation. ^31^ However, the potential impact of these crowders on the monomeric form of *α*S that determines its role in function or aggregation remains to be ascertained.

The intrinsic propensity of *α*S monomer to adopt variable helical and *β*-sheet rich fibrillar states in response to environmental cues, as reported in earlier studies, may have ramifications related to its biological function and toxicity leading to disease. This necessitates a comprehensive description of the structural and dynamical nature of *α*S monomer conformations in aqueous as well as in crowded media for understanding their roles in either preventing or triggering early stages of aggregation. Furthermore, it is of crucial importance to identify the key metastable states from the conformational ensemble of *α*S, and to ascertain the timescales of conformational transition for deducing the kinetics of downstream events leading to early self-assembly. With distinct amino acid composition, *α*S monomer comprises of (i) a basic N-terminal region (residues 1-60), (ii) a central hydrophobic region (residues 61 to 95) also known as non-amyloid-*β* component (NAC) region and (iii) an acidic C-terminal region (residues 96 to 140). The N-terminal region consists of a highly conserved hexamer motif (KTKEGV), shows relative helical propensity and can be considered as part of two helices, H1 (residues 1-30) and H2 (residues 30 to 60) when bound to micelles.^33–35^ The NAC region is highly amyloidogenic and identified as the critical determinant of *α*S fibrillation.^36^ However, the associated conformational heterogeneity of monomeric *α*S has precluded a clear consensus on its structure.

The various states adopted by *α*S are defined by molecular interactions that bear signatures of their propensity to prevent or enhance self assembly to form oligomers and eventually *α*S fibrils.^35, 37–40^ It has therefore stimulated research probing the structural and dynamical characterization of *α*S monomers as precursors of amyloid formation. A wide range of experimental studies were aimed at analysing the conformational features of *α*S monomer.^37, 39, 41–43^ Numerous studies have also used a combination of computational methods such as molecular dynamics and experiments to generate *α*S ensembles that are more attuned to experimental measurements.^35, 37, 38, 40, 44^ These studies have provided critical insights about the accessible states of *α*S in solution consisting of a heterogenous ensemble of compact to extended conformations. They revealed the presence of long-range tertiary interactions between the negatively charged C-terminal tail and the hydrophobic NAC or N-terminal residues; these conformations are potentially autoinhibitory towards oligomerization and aggregation. Moreover, single molecule experiments^45, 46^ and molecular simulations^47–49^ have also investigated the conformational dynamics of monomeric *α*S and measured the timescales of secondary and tertiary structure formation relevant to the pathological *α*S fibrils.

Many of the precedent reports investigating the conformation of *α*S monomer have made use of the atomistic precision rendered by variants of computer simulation and modelling. In majority of the investigations, the researchers have been prompted to take recourse of mixing MD simulation with numerous experimental techniques of different resolution via employing restraints or Bayesian approximation.^35, 37, 38, 40, 44, 49^ On the contrary, the bottleneck associated with accessing the time-scale required to sample heterogenous landscape of a long-sequence IDP such as *α*S via computer simulation has hindered its solo usage. In the present work we utilise sub-millisecond (0.108 millisecond) long MD simulation data of monomeric *α*S, generated at atomistic precision using a recently developed forcefield^50^ in explicit presence of water to characterise the key conformations that are mutually distinct. The extent of simulation data, to the best of our knowledge, is by far the longest reported for *α*S and is representative of an extensive, heterogenous ensemble. We utilise the framework of Markov state models (MSM)^51–53^ to dissect the metastable states and to evaluate the kinetics of conformational transitions between these states. Assorted based on the degree of compaction, we identify three macrostates namely, random-coil like extended state, intermediate compact state and globular-protein like compact or collapsed state. The intermediate compact state is the most populated state, with radius of gyration (*R_g_*) commensurate with previous experimental studies, followed by the globular-protein like state and lastly random-coil like extended state that is sparsely populated. These three macrostates can interconvert with transition times ranging from ∼0.1 *µ*s to ∼150 *µ*s and the fastest transitions lead to the dominant macrostate of intermediate compaction. We characterize these macrostates in terms of the intramolecular interactions and secondary structure that either stabilise the monomer or make them prone to aggregation, as shown by previous studies. Considering the increasingly appreciated effects of crowding on protein stability and self-assembly including the phase separation phenomenon, we further performed simulations of each of these metastable conformations in the presence of inert crowder molecules. The crowded environment induces variable degrees of compaction uncorrelated with the initial protein dimensions. The compaction is manifested in local stabilisation of the interactions and secondary structures present in the non-crowded ensemble as well as formation of new intra-and interdomain interactions. From comparison of *α*S monomer ensemble formed in non-crowded and crowded environments, we infer that crowding can modulate *α*S monomeric conformations both by harbouring as well as reducing the occurrence of aggregation-prone states. Finally, this work provides a comprehensive description of the structure and conformational transitions of potential metastable states of *α*S in solution and the impact of crowding on these states with relevance to favourability in the self-assembly pathway.

## Results and Discussion

### (I) Kinetic characterization of the conformational landscape of *α*- Synuclein

We started our investigation with a 73 microsecond-long continuous MD simulation trajectory generated at D. E. Shaw research using special-purpose supercomputer Anton2^54^ as the base ensemble of data in this investigation. In essence, the simulation was initiated from an extended conformation of *α*S using the recently developed a99SB-*disp* force field^50^ in explicit water and sampled at room temperature (see method for details). We first assessed the quality of the simulation data against NMR experiments and CD data (See Figure 1).We compared the secondary structure content of the ensemble with experimental estimates obtained from CD and chemical shifts. For the simulation ensemble, we used the STRIDE secondary structure assignment algorithm^55^ and VMD^56^ to calculate the residue-wise propensities to adopt *α*-helix and *β*-sheet states. The ensemble average secondary structure for helical and *β*-sheet content are 2.051 (± 2.926) and 6.709 (± 4.744) %, respectively. This is consistent with CD spectroscopic observations of 2% helix and 11% *β*-sheet content, respectively.^57^ We also used the *δ*2d program^58^ to calculate populations of secondary structure elements at the residue-level from backbone chemical shifts. As shown in Figure 1, there is a qualitative agreement between the experimental and simulation residue propensities. Importantly, we note that *β*-sheet propensity is peaked between residues 50 to 90 in the simulated ensemble that mainly covers the NAC region. The N and C-terminal regions also exhibit propensity to form extended structures, albeit lower than the NAC region. There is small *α*-helical propensity scattered across the N-terminal (residue 1 to 20) and NAC region (residues 55 to 65 and 85 to 95).

**Figure 1:**
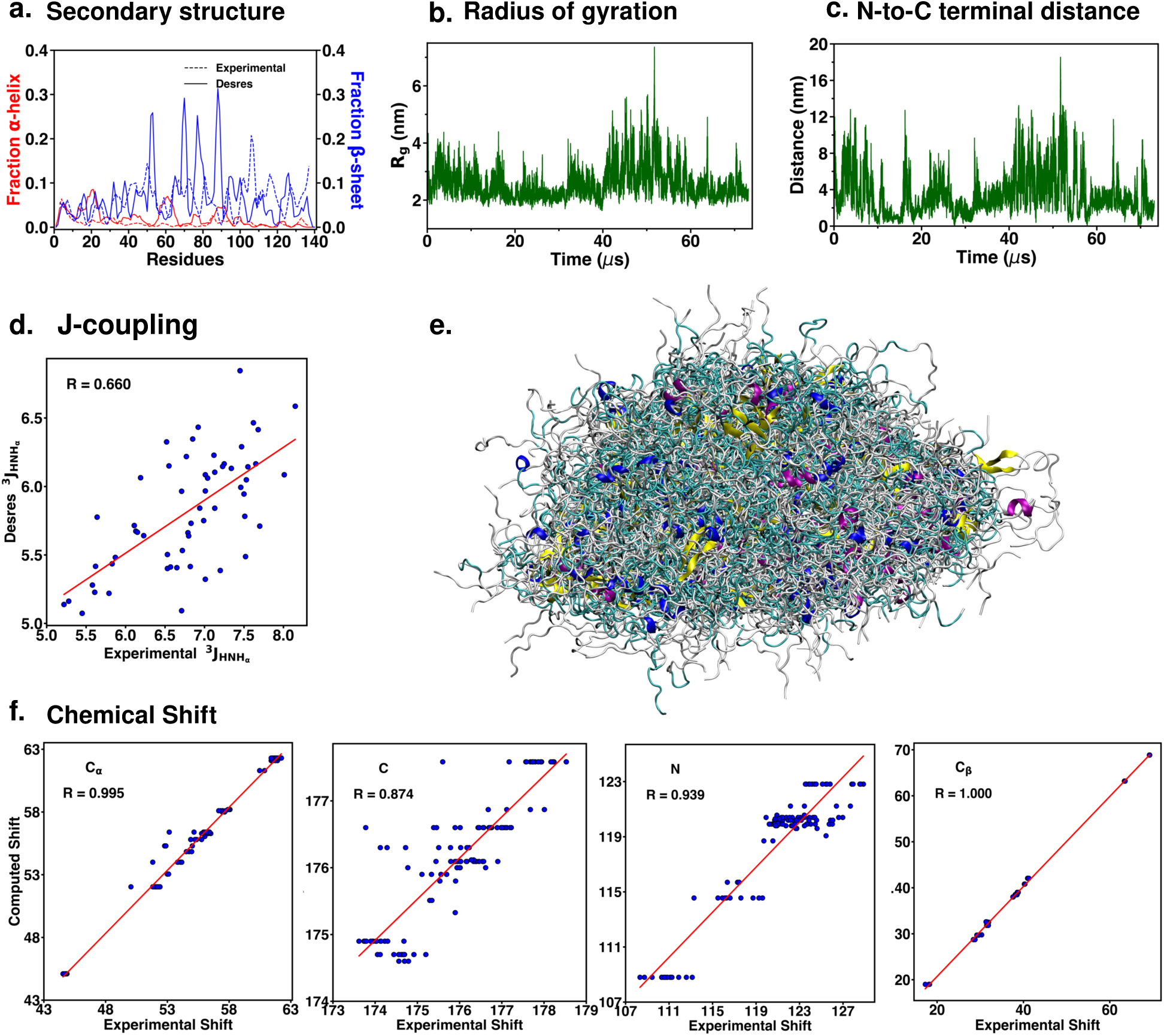
Heterogeneity captured in the 73 *µ*s-long *α*S simulation trajectory and validation of the simulation data against experimental data. (a) Fraction of *α*-helix and *β*-sheet propensities calculated from the simulation trajectory and experimental estimated from the program *δ*2d. Time evolution of (b) radius of gyration *R_g_* and (c) N- to C-terminal distances in the simulation ensemble. (d) Simulated and experimental ^3^*J_HNH_* scalar coupling. (e) Alignment of multiple structures within the *α*S ensemble. (f) Simulated and experimental chemical shifts of C*_α_*, C, N and C*_β_* atoms

Furthermore, as elucidated by the time-series plot of the radius of gyration (*R_g_*) of the protein, *α*S is found to sample a wide range of conformers of variable compaction between 1.5 nm to 7 nm. Interestingly, it has been reported that the average *R_g_* for a globular folded protein of 140 amino-acids (same amino-acid number as *α*S) is ∼1.5 nm, while the average *R_g_* for a random coil consisting of the same number of amino acids is ∼5.2 nm.^11, 35^ Similarly, the distance between N-terminal domain and C-terminal domain also represents a wide distribution suggestive of a heterogeneous distribution of long-range and short-range tertiary contacts. This indicates that the *α*S ensemble covers a broad conformational space occupying compact globular protein-like conformations to extended random coil-like states. The overlay of all *α*S conformations derived from the 73 *µ*s long MD trajectory depicts the conformational heterogeneity underlying the ensemble. These observations demonstrate the varying extent of secondary structure elements adopted by the conformations comprising the *α*S ensemble.

We also computed the chemical shifts of the *α*-carbons (C*_α_*), carbonyl carbons (C), amide nitrogens (N) and *β*-carbons (C*_β_*) using the SPARTA+ program.^59^ The chemical shifts calculated from the simulated ensemble shows excellent agreement with the corresponding experimental shifts indicated by high Pearson correlation coefficients of 0.995 (C*_α_*), 0.874 (C), 0.939 (N) and 1.0 (C*_β_*).^60^ In addition to chemical shifts, J-coupling constants provide residue-specific distributions of the backbone dihedral angles and therefore used to infer secondary structure. We measured the coupling of H*_N_* and H*α* (^3^*J_HNHα_* ) that is associated with the backbone torsion angle, *φ*, and estimated the correlation with NMR experiments. ^61^ It displays a reasonable agreement with experiments indicated by Pearson correlation coefficient of 0.6. However, it must be noted that J-couplings are unable to distinguish between unstructured peptides from their structured counterparts and hence are not the best observables for validating IDP ensembles. ^62^ These preliminary analyses attest to high quality of the *α*S structural ensemble captured by the underlying force field and sampling time. More crucially, these analyses also point out that the ensemble used to initiate the study is highly heterogeneous justifying the intrinsically disordered nature of *α*S.

The extent of structural and dynamical heterogeneity encoded in the 73 *µ*s MD simulation trajectory of monomeric *α*S (Figure 1) indicates that a quantitative kinetic clustering of this big-data is essential to identify the key metastable states underlying this complex ensemble. Towards this end, we undertook an initiative to build a comprehensive Markov State Model (MSM)^51–53^ of monomeric *α*S to ascertain key macrostates of finite life-time and elucidate the equilibrium of possible dynamical interconversions among them. A schematic of the framework implemented in the present investigation is depicted in Figure 2. The parent 73 *µ*s long trajectory served as the guide to spawn numerous short trajectories from structures extracted from different timepoints in an adaptive manner. For this purpose, this long trajectory was first discretized into five large clusters via *k* -means clustering algorithm^63^ using the radius of gyration (*R_g_*) and RMSD of the ensemble as the metric (see Figure 2); ten conformations from each cluster were then used to initiate new short simulations thus capturing a range of collapsed to extended states adopted by *α*S and the process was iterated to replenish the data-poor regime of the ensemble. The final ensemble, amounting to 108 *µ*s or 0.108 millisecond, was then used to build the MSM, which to the best of our knowledge, is the longest simulation dataset of monomeric *α*S.

**Figure 2:**
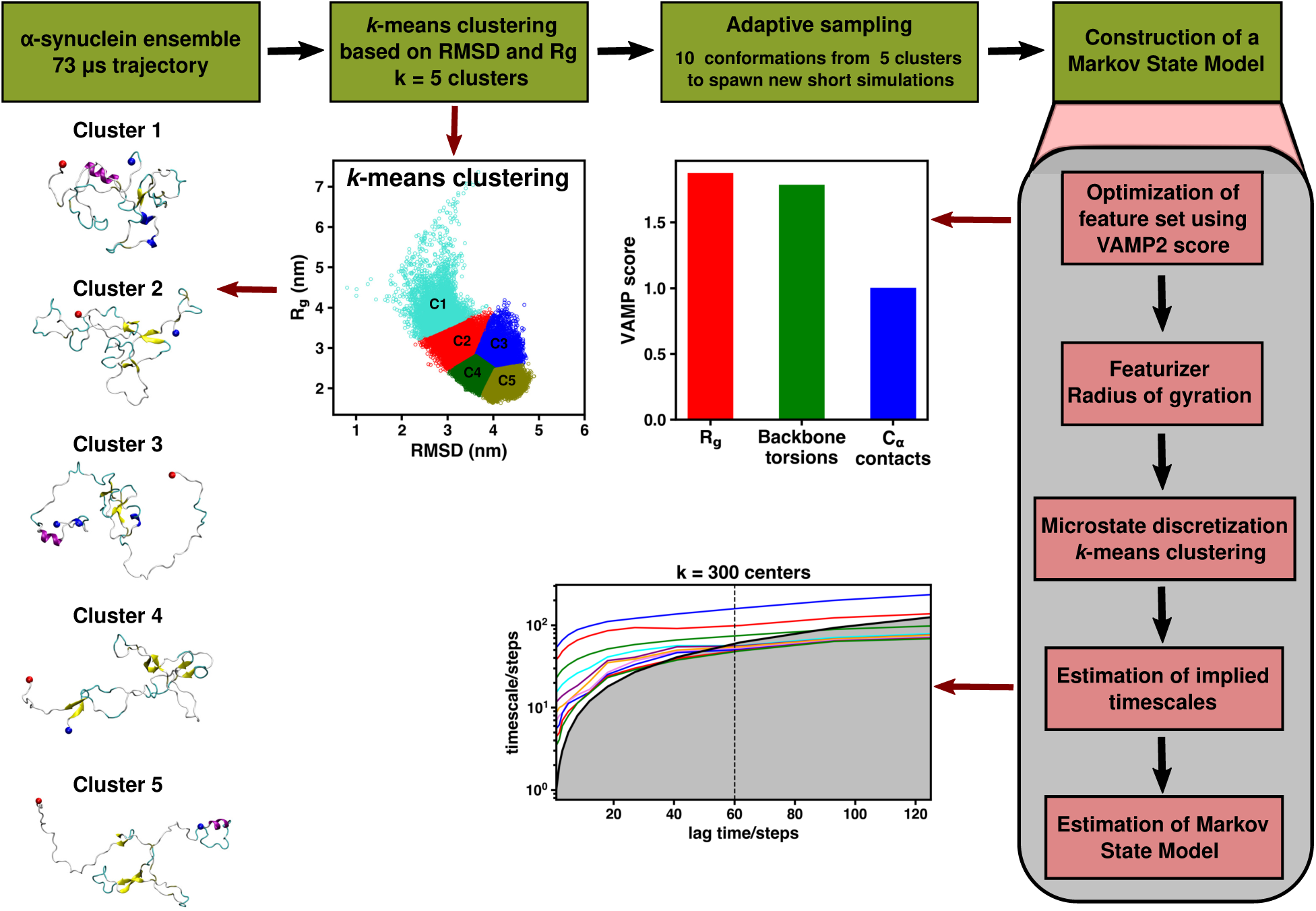
Illustration of the implemented workflow. A 2-dimensional scatter plot of the *k* -means clustering of RMSD and R*_g_* data. Representative snapshots of the five clusters are shown. VAMP-2 score ^64, 65^ estimated for the features *R_g_*, backbone torsions and C*α* contacts. Implied time scales (ITS) of underlying model; the dotted line marked at 60 ns is the MSM lag time chosen.

In the MSM protocol, the large ensemble of sub-millisecond long MD frames needs to be discretised into a large number of fine-grained microstates. To choose an appropriate discretisation metric for this complex IDP, we compared the optimality of a set of candidate features by comparing their respective VAMP-2 score. ^64, 65^ Recently introduced by Noé and co-workers, the VAMP-2 score is based on a variational approach for Markov processes (VAMP) that allows one to find optimal features, to cross-validate the hyper-parameters and to optimize a Markovian model of the dynamics from the given time series data.^64, 65^ The comparison of VAMP-2 score for three features of monomeric *α*S, namely, radius of gyration (*R_g_*), C*_α_* contact matrices and backbone torsions of the ensemble confirmed that *R_g_* shows the highest VAMP-2 score and hence would be most optimal for discretisation of the *α*S MD data. This also justifies the heavy usage of *R_g_* for describing the ensemble of monomeric *α*S or IDPs in general that lack a well-defined native structure and explore a diverse range of conformations over a wide range of sizes^35, 37, 49, 66^ Accordingly, we discretised the MD ensemble using *R_g_* and built a 300-microstate MSM.

The free energy profile, as derived from the MSM (see Figure S1) indicated a wide coverage of the ensemble along *R_g_*, with the basin located at an intermediate *R_g_* of 2.2 nm. The analysis of implied timescales plot (see methods for details and Figure 2) and Chapman-Kolmogorov (CK) test underlying the MSM (Figure S2) prompted us to kinetically cluster the 300-microstate MSM into three distinct macrostates (Figure 2). These three kinetically discrete states are identified as (i) extended state (MS1), (ii) compact intermediate (MS2) and (iii) collapsed state (MS3); multiple representative conformations of each of these macrostates is depicted in Figure 3. The mean *R_g_* values of these states are 5.85 (±0.43) nm, 2.59 (±0.45) nm and 1.95 (±0.08) nm, respectively; the probability distributions of these macrostates along *R_g_* are shown in Figure S1. The MSM is dominated by MS2 with a population of 85% while MS3 and MS1 populate 14% and 0.06% of the ensemble, respectively. The *R_g_* of the dominant state, MS2, is consistent with PRE-NMR measurements^37^ that reported an ensemble average value of 2.47 nm, ascertaining the prevalent views^35, 37^ that native monomeric *α*S is more compact than an equivalent random coil but more extended than a globular protein of same number of amino acids. Interestingly, the *R_g_* of the other two less-populated macrostates grossly represent the sizes of an equivalent random coil like polymer (MS1) and a globular protein (MS3). This is evident from the overlay of multiple conformations of each macrostate illustrated in Figure 3.

**Figure 3:**
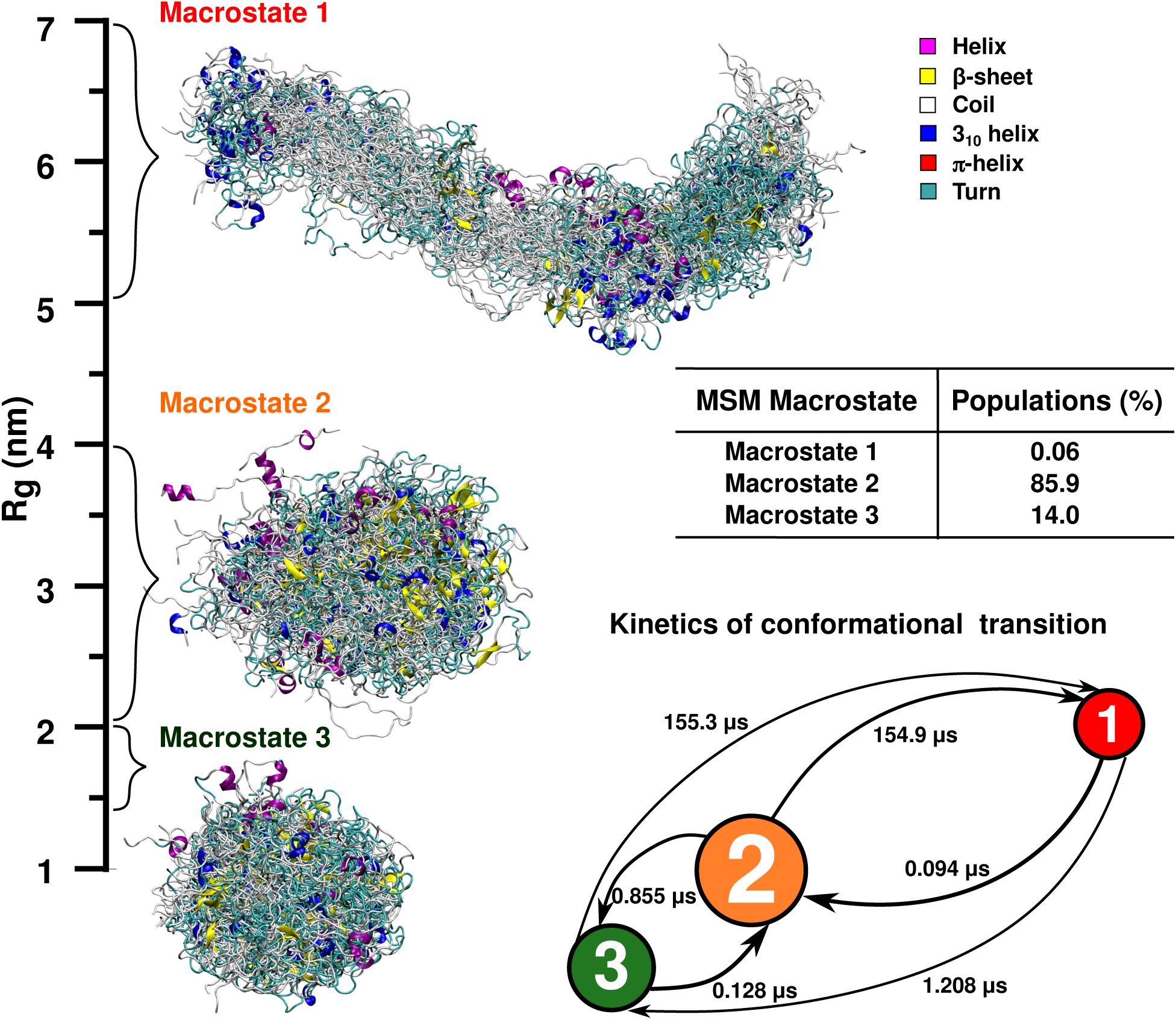
Overlay of conformations (colored by secondary structure) of the three macrostates estimated by the MSM is presented along the *R_g_* scale. The color key for the secondary structures is shown on the right. The equilibrium populations of the macrostates are tabulated. The kinetics of interconversion of the MSM macrostates is depicted. The three macrostates are represented as circular discs with arrows connecting them and the mean free passage times of transitions written over the corresponding arrow.

An analysis of the mean transition time between any two pairs of the metastable states (Figure 3) further indicates that the native ensemble (MS2) of *α*S dynamically interconverts with two other non-native states representing the random coil-like and globular protein-like dimensions. The timescale of interconversion is albeit very diverse, covering a wide range from a few nanoseconds to sub-millisecond. In particular, the native state MS2 undergoes a very rapid dynamical interconversion with the most compact non-native metastable state MS3 at a timescale of 0.128-0.855 *µ*s. On the other hand, the analysis suggests that the slowest transitions requiring ∼155 *µ*s occur from the compact MS2 and MS3 to the random-coil like extended transient state MS1. Importantly, the fastest transitions, occurring in ∼0.1 *µ*s, involves MS1 and MS3 attaining the intermediate compact MS2 conformation; therefore its status as the native, most populated state is justified. While previous studies incorporating experimental and computational methods have generated *α*S ensembles, the current approach, via building an MSM with an otherwise heterogenous ensemble, has enabled us to identify the key macrostates of variable compaction in an unsupervised manner and characterise their modes of interconversion. The identification of the metastable states and further quantification of the timescales of their interconversion from the MSM suggest that while navigating a broad spectrum of conformations in solution, *α*S monomer is predominantly more compact with faster timescales of transition arising from both random coil-like and globular protein-like states. In the following sections, we characterize the tertiary contacts prevalent in these macrostates.

### (II) Structural features of the metastable species

After identifying the key macrostates of *α*S monomer from an MSM, we further proceeded to characterize the structural features of these states and compare them across the ensemble. As discussed in the previous section, the most populated conformational ensemble discovered in the current MSM framework corresponds to a conformation of intermediate compaction (*R_g_* = 2.59 (±0.45) nm). The inter-residue contact probability map (Figure 4d), averaged over all the conformations of MS2 ensemble, shows intermittent presence of both intra-domain and long-range contacts in an otherwise weakly inter-connected network. In particular, we observed relatively intense contacts within the NAC region (residue 61-95) near the centre diagonal of the map (highlighted by dashed boxes in Figure 4d). The presence of these contacts within the NAC region is manifested by a relatively high propensity (∼25%) of *β*-sheet formation in this region (Figure 4f), amidst coil-like structure in most of the other locations. A representative snapshot in Figure 4e displays such *β*-sheets in the conformation. Furthermore, MS2 ensemble exhibits a small propensity of long-range interactions (Figure 4d), between the negatively charged C-terminus and the positively charged N-terminal residues. Such long-range intramolecular electrostatic interactions in *α*S have previously been detected in several experimental and computational studies. ^35, 37, 39, 40, 44^ These long-range interactions pertain to *β*-sheets formed with ∼10% propensity in these regions.

**Figure 4:**
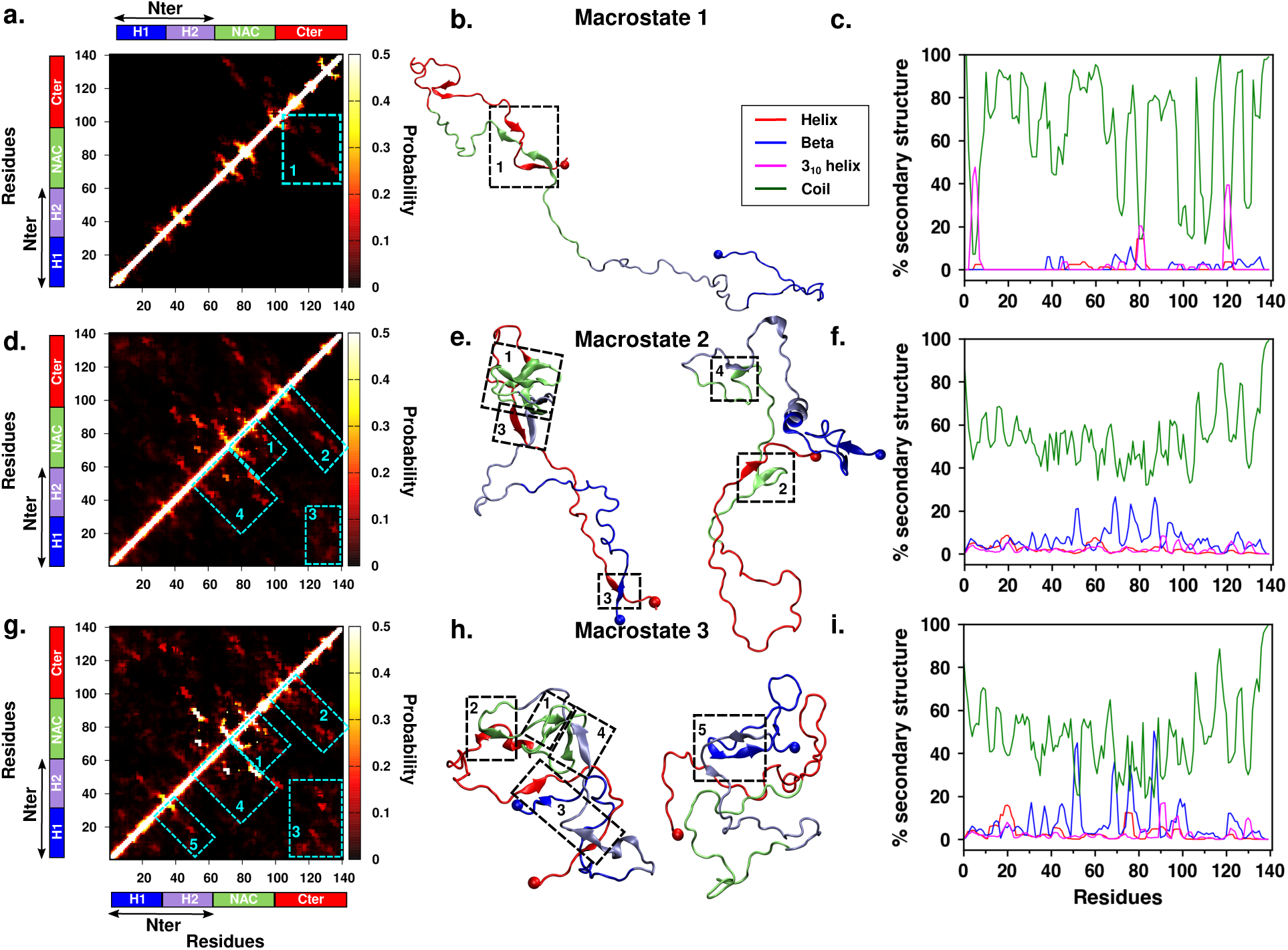
(a-c)Intrapeptide residue-wise contact probability maps of the three macrostates. Axes denote the residue numbers. The color scale for the contact probability is shown at the extreme right of each plot. The color bar at the top and left of each plot represents the segments in the *α*S monomer. (d-f) Representative conformations from each macrostate, colored segment-wise. (g-i) Residue-wise percentage secondary structure content of the helix (red), *β*-sheet (blue), 3_10_ helix (magenta) and coil (green) are shown for each macrostate

We compare the structural features of the most populated macrostate MS2 (Figure 4d-f) with the other two relatively less populated metastable ensembles, namely MS1 (Figure 4a-c) and MS3 (Figure 4g-i). A similar contact map analysis indicates that the most compact metastable ensemble MS3 reinforces many of the key aforementioned contacts discovered in MS2. In particular, this is characterised by higher probability of intra-domain contacts within the NAC region and relatively more enhanced long-range contacts between the N-terminal domain and C-terminal domain in the collapsed state MS3 (Figure 4g) than that in native MS2 (Figure 4d). These augmented interactions induce higher propensity of *β*-structures (∼40%) in MS3 and results in a compaction resembling that of a globular fold. On the other hand, the least populated extended conformations of MS1, while mostly in the coil state, display a low propensity for forming intramolecular contacts between the C-terminal and the central NAC region (see Figure 4a), that can be attributed to the transient formation of *β*-sheets while the N-terminus is mainly solvent-exposed. The secondary structure analysis of MS1 (Figure 4c) also indicates significant 3_10_ helicity in short segments of the N-terminal, NAC and C-terminal regions. However, the rapid kinetic interconversion between more populated sub-ensembles MS2 and MS3, as predicted by the MSM (see Figure 3) also indicates that the structural features of the overall ensemble will largely be dominated by the specific short-range NAC interactions and long-range interactions between the two termini.

### (III) Crowding-induced response of monomeric *α*-Synuclein

It is well appreciated that the biological milieu is extremely crowded and has profound impact on protein structure, dynamics and self-assembly.^22–25^ Experiments, therefore, are designed to mimic the intracellular crowding while studying protein stability, ligand binding, aggregation, etc. A wide range of crowders are used in experiments such as polyethylene glycol (PEG), polysaccharides like dextran and Ficoll, and other proteins such as lysozyme and bovine serum albumin.^26, 28, 29^ The effect of macromolecular crowding on IDPs is diverse relative to that of globular proteins.^67–69^ Furthermore, there is accumulating evidence on the nontrivial effects of crowding on the phenomenon of LLPS of IDPs.^31, 32^ A recent series of studies^31, 32^ on the phase separation of *α*S presents in vitro experiments to probe *α*S phase separation in the presence of the polymer crowder, polyethylene glycol (PEG). Importantly, evidences indicate that crowded conditions in vitro accelerates *α*S aggregation.^26–30^ This prompted us to evaluate the effect of crowding on the metastable states of *α*S monomer that we had derived via the MSM. We performed simulations of two representative conformations from each of the three macrostates (see Figure S3) in the presence of an inert spherical crowder. With 30% crowder volume fraction, we observe variable degrees of compaction of the three states in the crowded environment. The temporal evolution of *R_g_* and representative snapshots at the end of the simulation are displayed in Figure 5. We note that the extent of compaction is highest for the intermediate compact state MS2 with 37 to 46% compaction; the most extended conformation compresses by 12 to 26% while the most compact MS3 shows the least compaction of about 9 to 12%. These are consistent with the nonmonotonic trend of compaction in crowded environments reported in previous studies. ^68, 70, 71^ Based on a coarse-grained polymer IDP model spanning a wide range of polymer scaling regimes in the presence of purely repulsive spherical crowders, Miller et al. observed a non-monotonic relationship between the chain dimensions and the degree of compaction.^68^ They also showed that the free volume theory with a simple scaling factor accounting for the penetrability of expanded chains was able to quantitively reproduce the simulation results.

**Figure 5:**
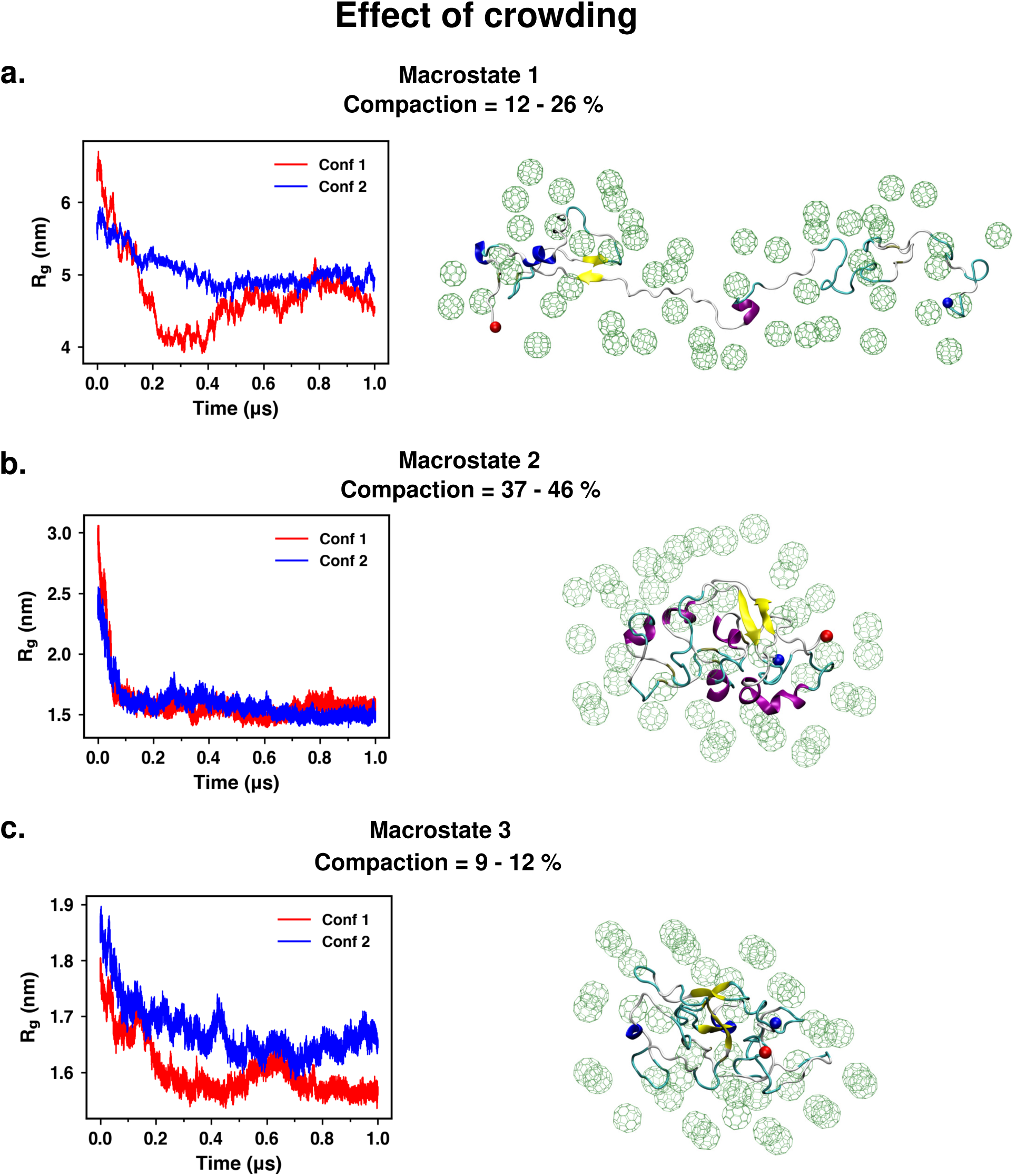
Evolution of the protein radius of gyration (R*_g_*) over the simulation timescale. A representative structure from the end of the simulation is shown. The peptide is colored according to secondary structure. The inert crowders in a shell of 0.8 nm are rendered as lines and colored in green

To understand the local structural changes induced by crowding, we analysed the intramolecular contacts and secondary structure content in the compact ensemble derived in the presence of crowders. Based on the time evolution of *R_g_* (Figure 5), we combined the last 500 ns from each of the two simulations performed for each macrostate as the ensemble formed due to the impact of crowding. In Figure 6, we present comparative analyses of residue-wise contact probability map and secondary structure propensities of the dominant macrostate MS2 in the presence and absence of crowders; representative structures from the crowding simulations are also depicted. It is evident from the contact map that many of the precedent off-diagonal contacts present in MS2 in non-crowded conditions (see upper half of the diagonal of contact map, figure 6 a) are amplified in presence of crowders (see lower half of the diagonal of contact map, figure 6 b). In addition, the presence of crowders also introduces several new inter-residue contacts and long-range interactions, which justifies the observation of maximal compaction of this macrostate upon crowding.

**Figure 6:**
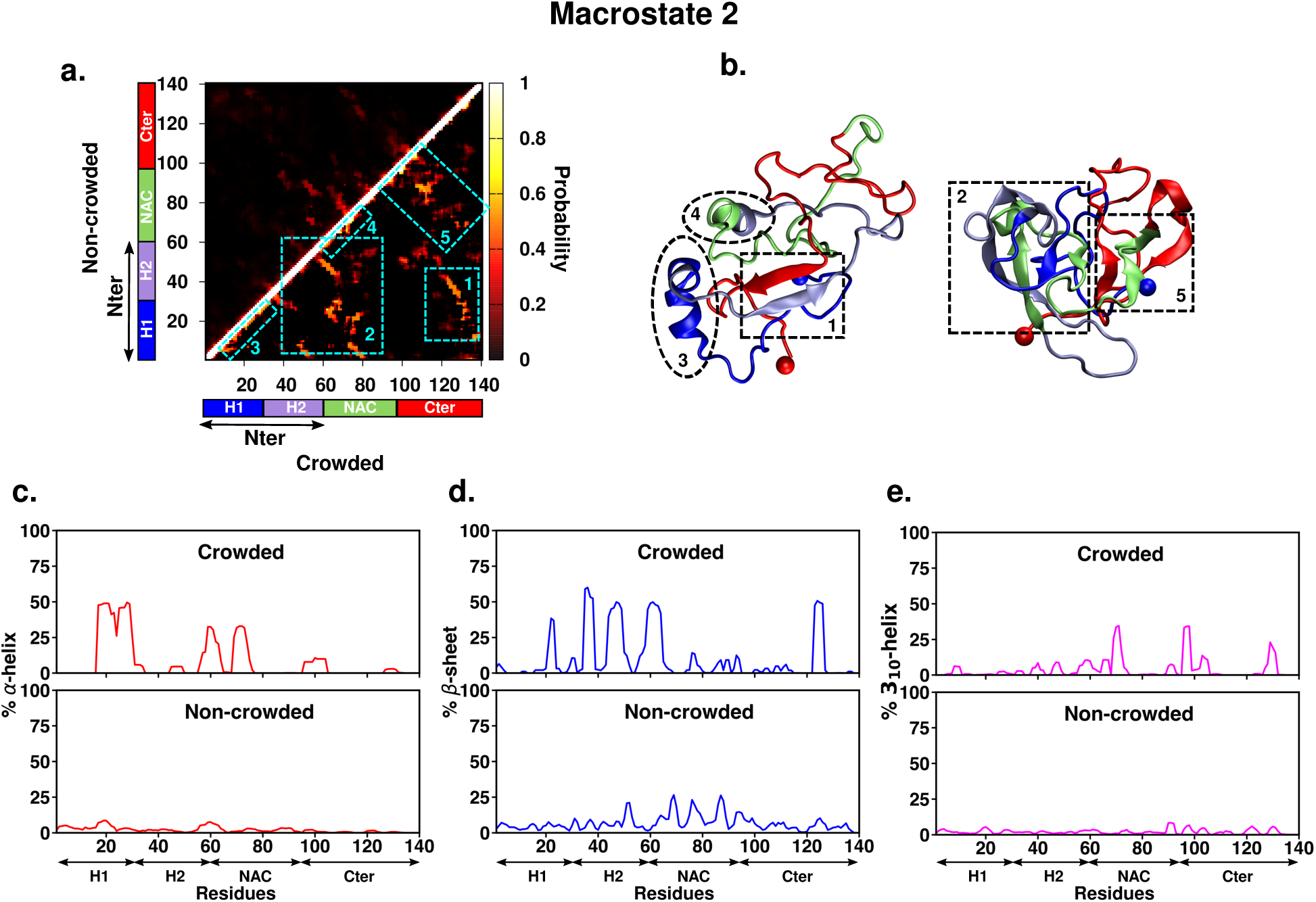
(a)Intrapeptide residue-wise contact probability maps of macrostate MS2 in the absence (upper half of diagonal) and presence of crowders (lower half of diagonal). The contact probability is indicated by the color scale on the right of the plot (b) Representative conformations from crowding simulation ensemble are shown and colored segment-wise. Dashed boxes in cyan are marked in the crowded system contact map to indicate key contacts and the corresponding secondary structures are marked and numbered in the representative structures. Percentage secondary structure propensity of each residue in the crowded (upper-panel) and non-crowded (lower-panel) systems are presented for (c) *α*-helix (d) *β*-sheet and (e) 3_10_ helix

The conformational features described below can be visualized in the representative snapshots and are annotated based on the contact map. The collapsed C-terminal region establishes long-range interactions with both N-terminal and NAC regions. Specifically, the end segment of the C-terminal consisting of residues 123 to 139 forms contacts with residues 1-30 of the N-terminal; these contacts are also formed sparsely in the absence of crowders. On the other hand, crowding-induced compaction gives rise to high density of new contacts arising from the *β*-sheet formed between residues 100-122 of the C-terminal and residues 31-60 i.e. H2 segment of the N-terminal region. The hydrophobic NAC region is also involved in long-range interactions with most of the C-terminal region in the crowded conditions. These regions also interact weakly in non-crowded conditions however via different contacts. The intra-domain interactions forming the *β*-sheet network in the NAC region as observed in the non-crowded condition are evidently reconfigured due to the impact of crowding (see the marked region in Figure 6). The NAC region in the compact ensemble, by contrast, forms helical structures, both *α*-helix and 3_10_ helix, with ∼35 % propensity. The ensemble also comprises of conformations in which the NAC region and N-terminal region form *β*-sheet network with propensities ranging from ∼10-50 %. Finally, the N-terminal region from residues 16-32 form *α*-helices in a significant population of the MS2 compact ensemble. These aforementioned interaction patterns displayed by the charged N-terminal region are practically absent in the MS2 ensemble in the non-crowded system, and hence can be considered to be driven by compaction in crowded systems. Also, short residue patches exhibit small tendency (∼10-30 %) to form 3_10_ helices and such patches are spread across the entire protein sequence. We however, note that a small fraction of these observed interactions are reminiscent of the initial conformations used for simulations in the presence of crowders (see Figure S3) and are further amplified along with formation of new contacts. The extent of structural rearrangements observed in MS2 ensemble in the presence of crowders corroborates with the maximal compaction attained by this state in crowded environments compared to the other two macrostates.

Similar comparisons of the contact probabilities relevant to the secondary structural changes and representative snapshots are provided for the other two macrostates MS3 and MS1 in Figure S4 and Figure 7, respectively. The globular-protein like MS3, which is least affected by crowding, does not display major differences in the nature of interactions but are present with much higher propensity compared to the non-crowded ensemble. The NAC region and part of the N-terminal H2 segment comprising of residues 50 to 85 form an extensive *β*-sheet network with ∼75-100 % propensity. As seen in the non-crowded ensemble, there are N-to-C long-range interactions (residues 1 to 10 and 130 to 135) manifested as *β*-sheets in ∼50 % of the ensemble. The comparison of MS1 ensemble in the absence and presence of crowders indicates that moderate compaction caused by crowding elevated the existing contacts to an appreciable extent along with development of a few new features in this macrostate (see Figure 7). As seen in the non-crowded system, while the N-terminal remains extended and solvent exposed, the C-terminal region folds to form *β*-sheet network with the NAC region (residues 70 to 95) in 15 to 30 % of the ensemble. Also, in 10-30 % population of the structures, the C-terminal residues 110 to 130 adopt 3_10_ helicity. A notable new feature observed in the crowded MS1 ensemble is the appearance of *α*-helical nature extending from end of H2 region to a short segment marking the beginning of the NAC region (residues 50 to 70) that is present in nearly 50 % of the structures (see snapshot and secondary structure in figure 7). These analyses provide insights about the response of *α*S monomer to crowded conditions dictated by the protein dimensions as well as the effect of crowding on the structural features of these ensembles. In the upcoming section, we discuss the impact of the conformational features of *α*S macrostates in crowded and non-crowded conditions with relevance to their implications in the fibrillation pathway.

**Figure 7:**
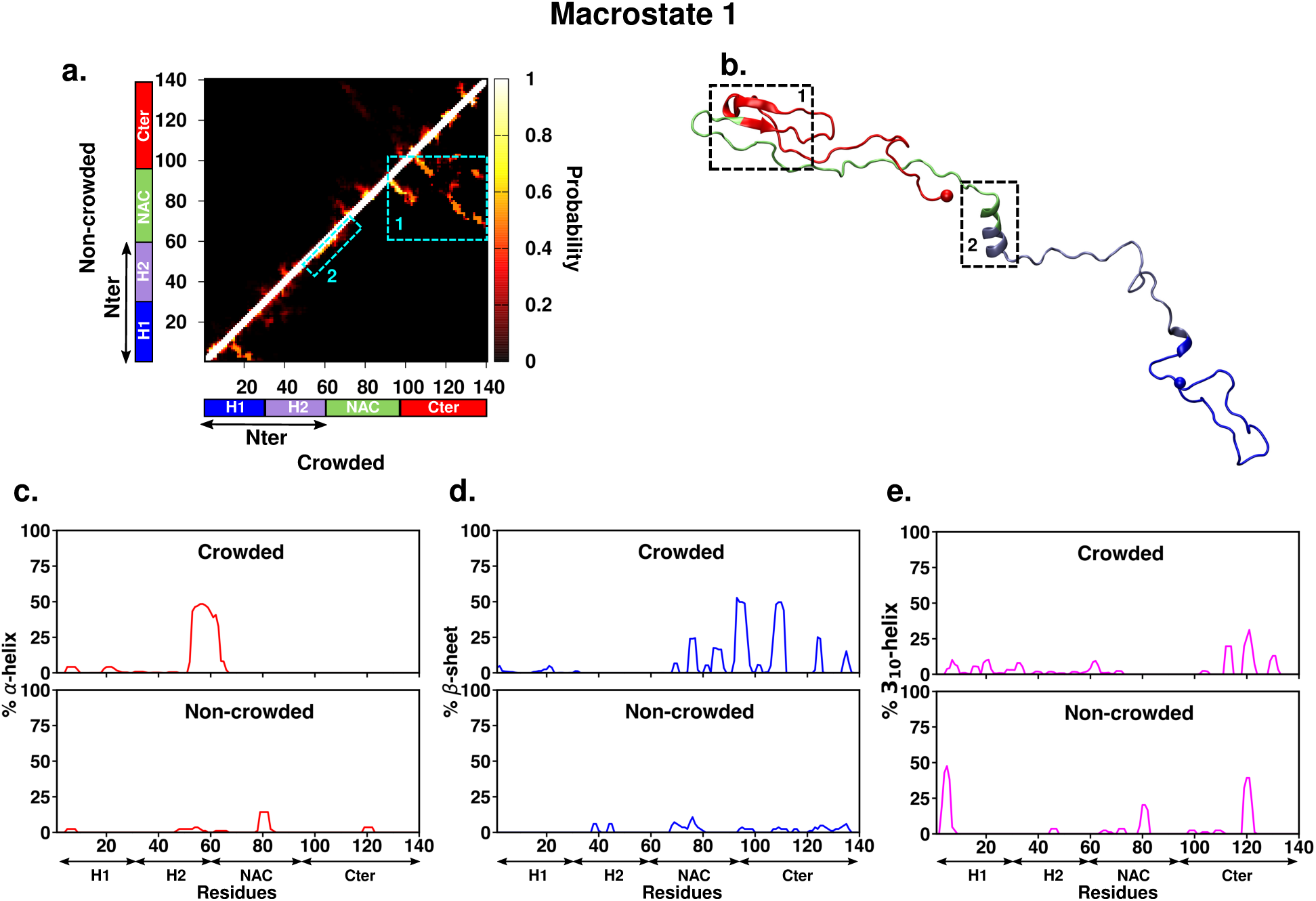
(a)Intra-peptide residue-wise contact probability maps of macrostate MS1 in the absence (upper half of diagonal) and presence of crowders (lower half of diagonal). The contact probability is indicated by the color scale to the right of the plot (b) Representative conformations from crowding simulation ensemble are shown and colored segment-wise. Dashed boxes in cyan are marked in the crowded system contact map to indicate key contacts and the corresponding secondary structures are marked and numbered in the representative structures. Percentage secondary structure propensity of each residue in the crowded (upper-panel) and non-crowded (lower-panel) systems are presented for (c) *α*-helix (d) *β*-sheet and (e) 3_10_ helix

### (IV) Monomer ensemble and implication on fibrilization/aggregation

Our aforementioned analyses indicate that the *α*S monomer can assume a plethora of conformational states by reconfiguring the molecular interactions and these changes are sensitive to the surrounding environment such as presence of inert crowders. Intensive research efforts have been directed towards distinguishing the functionally relevant conformations from the structural features that render the monomer more prone to aggregation.^37, 39–43, 47, 49, 72^ These studies have shown that while being largely unstructured in solution, residual secondary structure and intramolecular interactions are present in *α*S monomer. Experimental evidences have demonstrated that under native conditions, *α*S exists as stable *α*-helically folded tetramer that prevents it from self-aggregating. ^9, 10^ It has been therefore postulated that destabilization of these helical forms precede the misfolding and aggregation process. Long-range electrostatic interactions between the positively charged N-terminus and negatively charged C-terminus, as well as hydrophobic interactions between the C-terminal residues and NAC region have been reported in several previous computational and experimental studies done using a range of biophysical techniques.^35, 37, 39, 40, 43, 44, 49, 73^ The presence of such long-range interactions were proposed to sequester the hydrophobic NAC region from self-association and thus such autoinhibitory conformations serve to inhibit aggregation. The hydrophobic NAC region is the minimal segment required for in vitro *α*S fibrillation^74^ and in the aggregated forms, the fibril core consisting of residues 30 to 100, is mainly engaged in *β*-sheet interactions,^35, 47, 75–77^ which is the universal feature of amyloid fibrils. This indicates that the conformations rich in *β*-sheet interactions in these critical regions are predisposed to the aggregation phenomenon.

With this background on functional and potentially pathological role of conformations, we parsed our MSM-generated monomeric conformational ensembles and corresponding contact maps in a bid to curate the signatures of *aggregation-prone* or *aggregation resistant* interactions present among these MSM-extracted metastable states. Figure 8 illustrates the key interactions identified in the current non-crowded and crowded ensemble, demarcated on the basis of whether they contribute to aggregation-prone or aggregation-resistant conformations. In the non-crowded ensemble (Figure 8a), short segments spread across the entire N-terminal region are involved in electrostatic interactions with mainly the end of the C-terminal region. Hydrophobic interactions are also present between the NAC region and end of the C-terminal region. These are manifested as *β*-sheet interactions between these regions. Short stretches of residues in the N-terminal and NAC region adopt *α*-helical structure. Such interactions are signatures of conformations that are potentially resistant to forming intermolecular interactions with other peptides and thus resist aggregation. On the other hand, in some populations of the non-crowded ensemble, the NAC region and the H2 segment of the N-terminal region interact via *β*-sheet interactions; there are *β*-sheet interactions present in the N-terminal region. Such interactions enhance the susceptibility of *α*S monomer to self-assemble and form oligomers and higher-order aggregates.

**Figure 8:**
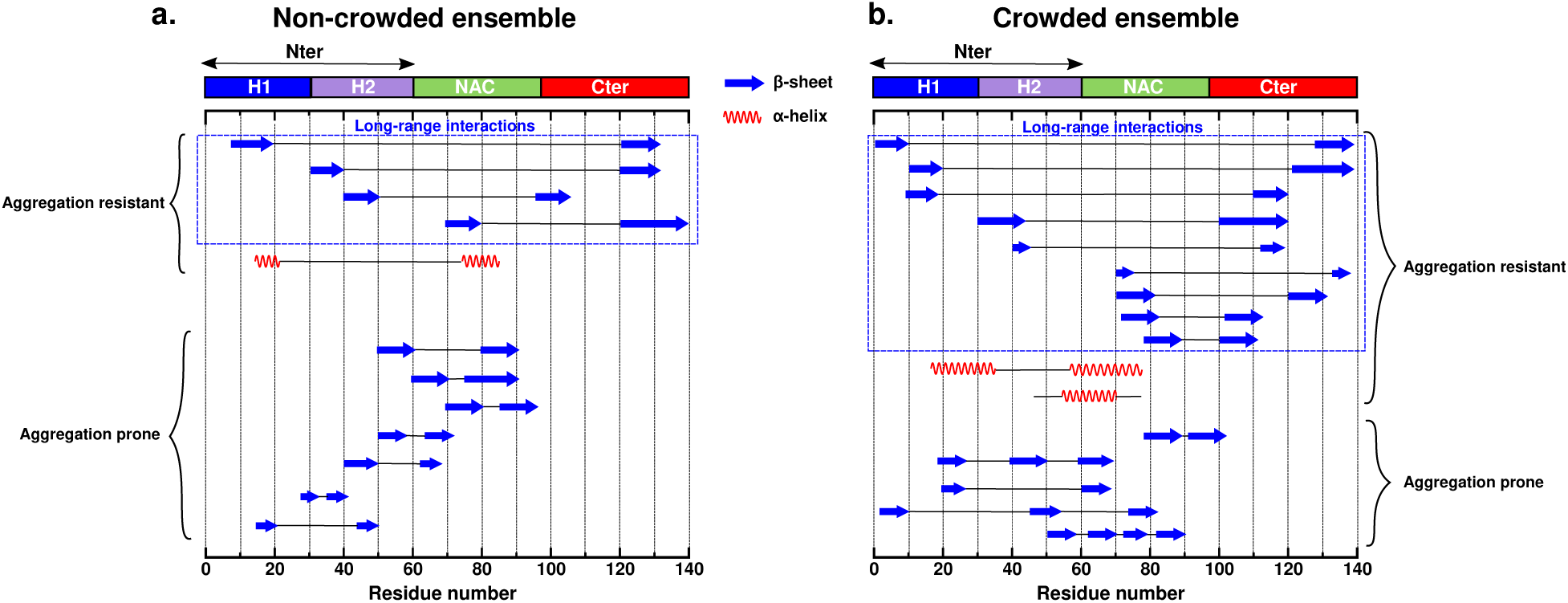
Schematic depicting the various intramolecular interactions in the *α*S ensemble assorted in the current work in a) non-crowded and b) crowded conditions. The interactions manifesting as secondary structures (*α*-helix and *β*-sheet) are shown. The interactions that are potentially aggregation-prone and aggregation-resistant are marked.

The interaction profile of the ensemble generated in crowded conditions (see Figure 8b) reflect the role of crowding in modulating the conformational preferences. Comparison with the non-crowded ensemble indicates that there is significant enhancement in the interactions that govern the aggregation resistance of these conformations. The crowding induced compaction is manifested in the C-terminal-NAC *β*-sheet interactions that is more prominent relative to the non-crowded ensemble. There is also a relative increase in the long-range tertiary contacts between the N-teminal and C-terminal region, since the compaction now causes the entire C-terminal region to form *β*-sheet network with the N-terminal region. These observations suggest that in a crowded environment, *α*S monomer preferentially favours long-range interactions to adjust to the excluded-volume effects of such environments. Furthermore, in some sub-populations of the ensemble, the NAC and N-terminal regions adopt *α*-helical nature that is reminiscent of the native-like helical species reported in earlier studies.^9, 10^ Thus, with increased propensity of such structural features, crowded conditions favoured the formation of aggregation-resistant species. Conversely, the ensemble also has subpopulations of conformations in which there is intensified network of *β*-sheet interactions within the NAC region as well as interdomain interactions between the N-terminal and NAC regions as compared to similar conformations in the non-crowded ensemble. Such conformational species may potentially stimulate oligomerization and fibrillation.

Overall, the above results depicted schematically presents the assortment of monomer *α*S conformations in non-crowded and crowded conditions. A comparative analysis of the two ensembles provides insights on how crowding can navigate them towards inhibiting or promoting aggregation. Essentially, *α*S monomer tends to populate a range of different conformations depending on the environment that can interconvert and follow different routes culminating in stable, functionally viable states or pathologically-relevant aggregated species.

## Conclusion

*α*S is categorized as intrinsically disordered and while being unstructured in solution, it can adopt conformations that vary in fundamental structural features such as the nature and extent of secondary structure as well as tertiary packing. Depending upon the nature of these interactions, some of the conformations may be suitable for its physiological functions while some conformations may be predisposed to aggregate with other partners into higher ordered species. The vast literature on *α*S indicates that functional state of this protein in predominantly helical^9, 10, 78–80^ whereas when it associates with other *α*S peptides, the aggregates are rich in *β*-sheet content.^11, 12^ These observations indicate that *α*S monomer ensemble is highly heterogenous and skewed as a function of the environmental conditions.

They underscore the importance of characterising *α*S monomer ensemble and determining the conformational features that can either trigger early events in aggregation or hinder the process. It is also critical to generate these ensembles in a crowded environment that mimics the cellular milieu since macromolecular crowding can significantly modulate the conformational space accessed by proteins relative to aqueous conditions. Multiple studies have demonstrated the acceleration of *α*S aggregation in crowded enviroments.^26–30^ Moreover, recent reports have also shown liquid-liquid phase separation of *α*S that is promoted in the presence of molecular crowders owing to increased local concentration.^31, 32^

The heterogenous and dynamic nature of *α*S ensemble pose challenges in its characterisation, both experimentally and computationally. However, numerous biophysical techniques such as NMR, small-angle X-ray scattering, CD and computational methods such as MD simulations have been used in a standalone manner or in combination to delineate the features of the monomeric state.^35, 37–43, 45–49^ These studies have provided key insights about the accessible states, their conformational features, intramolecular interactions and their potential role in self-assembly as well as timescales of secondary and tertiary structure formation. The overarching goal of our study is to characterise the metastable states of *α*S and assess the impact of crowded environments on these states to compare and delineate the two ensembles in terms of aggregation-prone and aggregation-resistant conformations inferred from existing literature.

Using the current state-of-the-art approach of complementing adaptive sampling with Markov State Models, we elucidate the metastable states of *α*S monomer in aqueous solution and the kinetics of their interconversion. Based on *R_g_* that was determined as the most optimal collective variable to build the MSM, the resultant model yielded three macrostates with mean *R_g_* values of 5.85 (±0.43) nm (MS1), 2.59 (±0.45) nm (MS2) and 1.95 (±0.08) nm (MS3). Notably, state MS2 is predominantly populated in the ensemble (85%) and its mean *R_g_* value corroborates PRE-NMR studies of *α*S monomer.^37^ This dominant state MS2 is significantly more compact than the random coil like state MS1 and relatively more expanded than the globular protein-like state MS3. We estimated the transition timescales among these macrostates, which revealed that the fastest transitions occur from the states MS1 and MS3 to the intermediate compact state MS2 within timescales of ∼0.1 *µ*s. On the other hand, the most compact MS3 interconverts with MS2 over ∼0.8 *µ*s while the slowest structural transition occurs during the expansion of the compact states MS2 and MS3 to the extended MS1 conformation. The kinetics of structural interconversion also further validates the preference of *α*S to attain the intermediate compact conformation in solution.

We further discerned the intramolecular interactions and secondary structure of the three metastable states. In the compact states MS2 and MS3, the oppositely charged N and C-termini interact via long-range interactions. Such long-range interactions manifest as *β*-sheet interactions among these regions. Such interactions have been observed in earlier studies on *α*S monomer and suggests that the presence of such long-range interactions can potentially shield the NAC region that makes it less susceptible to aggregation. ^36, 37, 39, 40, 44, 49^ All the three states also display minor propensity of long range interactions between the negatively charged C-terminal and the hydrophobic NAC region. In MS2 and MS3, the NAC region is also involved in *β*-sheet interactions that can promote aggregation-prone conformations since this segment is part of the fibril core and is present in *β*-sheet form in these assembled species.

On exposing the metastable states to crowded environments, they undergo compaction but the response is variable and non-monotonic. The dominant state MS2 gets compacted by a substantial amount in presence of inert crowders, followed by the extended state MS1, while the originally compact state MS3 is least affected. The compaction is also manifested as enhancement in the intramolecular contacts either by stabilizing the existing interactions or formation of new contacts. The long-range tertiary contacts between the terminal regions and those formed between NAC and C-teminal region are discernably enhanced in the crowded ensemble. Some subpopulations are also more helical in the N-terminal and NAC regions. Such interactions make the conformations more favorable to sustain in the monomer state, thus preventing aggregation. The ensemble also harbors conformations that are primed for oligomerisation via sticky interactions such as *β*-sheet network in the NAC and N-terminal regions.

The identification of conformations that can be potentially competent or incompetent to aggregate and pursue fibrillation is an important step towards rationalising the molecular events that trigger *α*S-associated pathologies. This knowledge can be leveraged to design therapeutic strategies against various synucleinopathies and small molecule discovery for targeting *α*S^81^ .

## Methodology

### Unbiased Adaptive Sampling Simulation

A 73 *µ*s-long continuous MD simulation trajectory of *α*S generated using Anton supercomputer by D. E. Shaw research and made accessible to the authors, was used as the starting ensemble for implementing adaptive sampling strategy in this study. The starting structure for this simulation is an extended conformation of *α*S modelled using the a99SB-*disp* force field and solvated using a99SB-*disp* water model^50^ . After equilibrating the system for 1 ns at 300 K and 1 bar using the Desmond software,^82^ production runs were performed at a time step of 2.5 femtoseconds, in the NPT ensemble with the Anton specialized hardware. ^54^ In the adaptive sampling approach, short MD simulations were run starting from a multitude of conformers obtained from the long parent trajectory. A flowchart of the protocol employed is presented in Figure 2. To extract starting conformations for these short independent runs, the 73 *µ*s long simulation trajectory was subjected to *k* -means clustering^63^ based on structural root mean squared deviation (RMSD) and radius of gyration (R*_g_*) of the ensemble. A 2D scatter plot of the five clusters is shown in Figure 2. Ten representative conformations from each of the five clusters are selected as starting structures for independent short simulations. These conformations span a wide range of dimensions from extended to compact conformations; the ensemble thus sampled encompasses the gamut of possible conformations adopted by *α*S in solution. The simulation details of these short simulations are described below. The protein was solvated by water molecules in a dodecahedron box of dimension 14.5 × 14.5 × 10.25 nm and periodic boundary condition was implemented in all three dimensions. The average number of water molecules in the box was 70600 and nine sodium ions were introduced to render the system electroneutral. The total number of particles in the system was ∼ 284500. The all-atom a99SB-*disp* force field and water model^50^ were used to model the protein and solvent. The unbiased MD simulations were performed using the GROMACS 2020 simulation package.^83, 84^ A time step of 2 fs and the leap-frog integrator was used. The simulation was performed in the isothermal-isobaric (NPT) ensemble at an average temperature of 300 K maintained using the v-rescale thermostat^85^ with a relaxation time of 0.1 ps and a pressure of 1 bar using the Parrinello-Rahman barostat^86^ with a time constant of 2 ps for coupling. The Verlet cutoff scheme ^87^ with 1.0 nm cutoff was applied for Lennard-Jones interactions and short-range electrostatic interactions. Long-range electrostatic interactions were computed using the Particle-Mesh Ewald (PME) summation method^88^ and covalent bonds involving hydrogen atoms were constrained using the LINCS algorithm.^89^ All the systems were minimized using the steepest descent algorithm, followed by equilibration in the isothermal-isochoric (NVT) and subsequently in the NPT ensemble with position restraints on the heavy atoms of the protein. A large number of adaptively spawned simulations with a saving frequency of 100 ps and variable timescales were performed, amounting to a cumulative length of ∼35 *µ*s . Together, the ensemble amounted to an aggregated (73 + 35) = 108 *µ*s of atomic resolution data.

### Construction of the Markov State Model

We employed PyEMMA,^90^ a Markov State Model (MSM) building and analysis package to identify the metastable states of *α*S and estimate the kinetics of their transitions using the aforementioned heterogenous ensemble of of ∼108 *µ*s trajectories (see Figure 1e for a representative image) . In order to build an MSM, the first step is to compute a kinetically-relevant distance metric or feature that describes the conformational ensemble.^53^ We selected the radius of gyration (*R_g_*) of the protein as the input feature. The feature selection was done based on VAMP-2 scores (Variational approach for Markov processes) ^64, 65^ that can be used to rank features by measuring the kinetic variance among the features. We considered three different featurizations for this estimation: (a) *R_g_*, (b) backbone torsions and (c) C*α* contact maps; the VAMP-2 scores for these features are shown in Figure 2. This analysis indicates that the highest scoring *R_g_* is the most suitable feature for the MSM. Next, the *k* - means clustering algorithm is applied to the *R_g_* feature space to discretize the conformations in the trajectories into 300 clusters or microstates. The transitions between the microstates are then counted to build a transition matrix, at a specific lag time at which the model is Markovian. In order to choose an appropriate lag time, we built the 300 microstate model over a range of lag times. The implied timescales (ITS) or relaxation timescales implied by the MSM at a particular lag time can be evaluated and the timescale at which the ITS plot levels off can be chosen as the appropriate lag time to build the model. Based on the ITS plot (see Figure 2), we chose a lag time of 60 ns to build the 300 microstate MSM. The clear gap between the second and third eigenvalues in the ITS plot suggests that there are three well-resolved metastable states in the underlying free enengy landscape of *α*S monomer. Thus, a three-state coarse-grained model was built at a lag time of 60 ns using the Perron Cluster Cluster Analysis (PCCA+) method. ^91^ A Chapman-Kolmogorov test ^51^ was used to check if the model is Markovian; the output of the test is depicted in Figure S2. Finally, the transition path theory^92–94^ was used to ascertain the transition paths and fluxes.

### Modelling crowded environments and MD simulation protocol

Two representative conformations from each of the three metastable states were selected to study the effect of crowded environments. We used C60 fullerene used in prior studies^95^ as the small molecule crowder to mimic crowded environment. The inertness of the crowder was modelled via introducing repulsive interactions with the protein atoms and other fullerene molecules. On the other hand, the interactions of the crowder with the solvent and ions were modelled using both attractive and repulsive interactions. We set up the crowding simulations with each selected *α*S conformation packed with a random distribution of the crowder molecules occupying 30% of the simulation box volume. This was followed by solvation and neutralizing the system by adding ions . The details of the simulation method are as mentioned in the previous subsection. The simulation details of the crowding simulations are provided in Table S1. The six crowding simulation trajectories were performed for 1 *µ*s each. The ensemble generated from the last 500 ns of each of these trajectories was used for further analyses of the crowding-induced compact states.

## Supplemental Information

All supplemental figures described in the article (PDF)

## Acknowledgements

We sincerely thank D. E. Shaw research for providing us with access to a long simulation trajectory of monomeric alpha-synuclein. We acknowledge support of the Department of Atomic Energy, Government of India, under Project Identification No. RTI 4007. JM acknowledges Core Research grants provided by the Department of Science and Technology (DST) of India (CRG/2019/001219).

