## Supplemental figures and table for "Conformational Ensemble of Monomeric *α*-Synuclein in Aqueous and Crowded Environments as revealed by Markov State Model"

### Supporting information for 'Conformational Ensemble of Monomeric $\alpha$-Synuclein in Aqueous and Crowded Environments as revealed by Markov State Model'

Sneha Menon and Jagannath Mondal*

*Tata Institute of Fundamental Research, Center for Interdisciplinary sciences, Hyderabad 500046, India*

,+914020203091

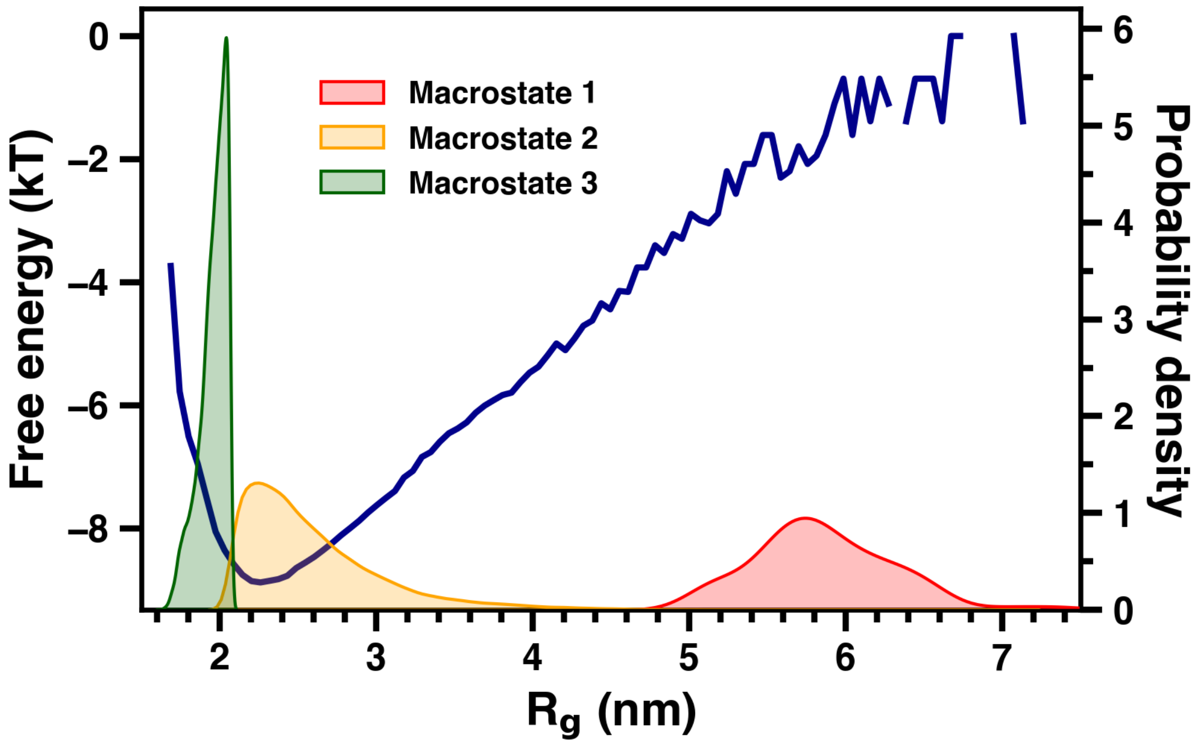


Figure S1: Probability density of the three macrostates overlaid with the one-dimensional free-energy landscape as a function of $R_g$.

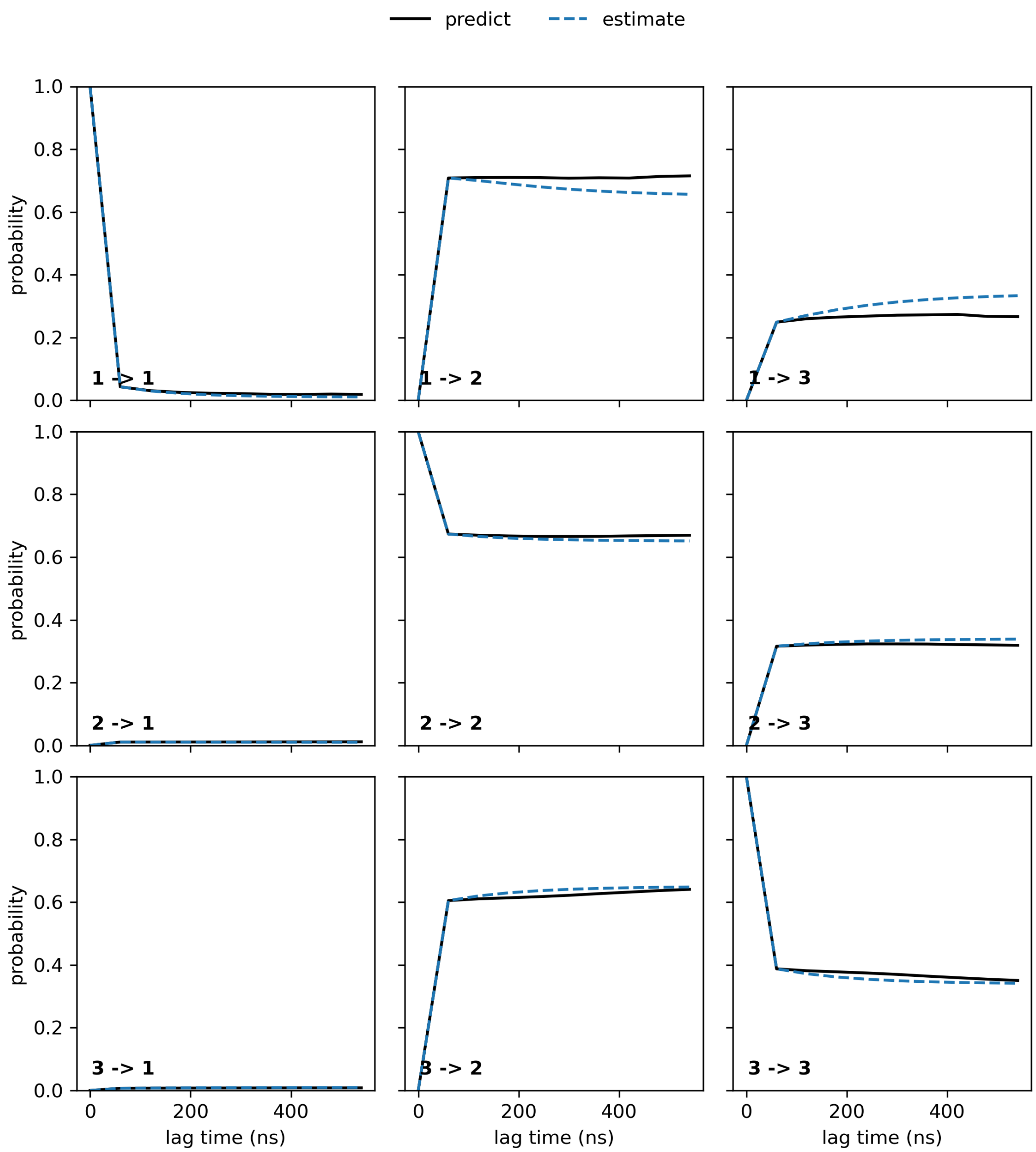


Figure S2: The Chapman-Kolmogorov test performed for the 3-state Markov State Model built at lag time 60 ns

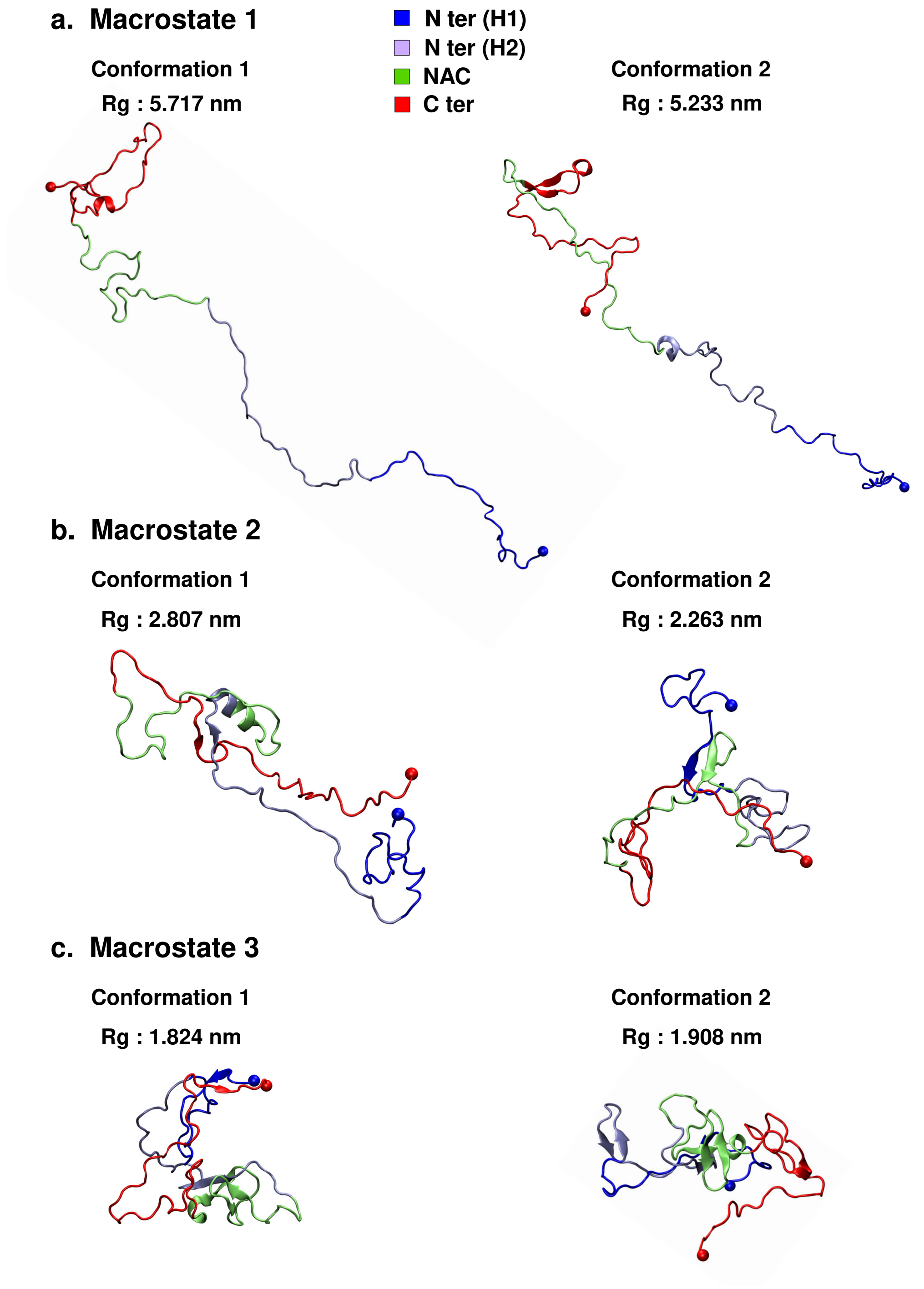


Figure S3: The initial conformations of $\alpha$S monomer used for simulations in the crowded environment. Two conformations for each macrostate (a) MS1, (b) MS2 and (c) MS3 are depicted. The structures are colored according to the regions defined in the main text.

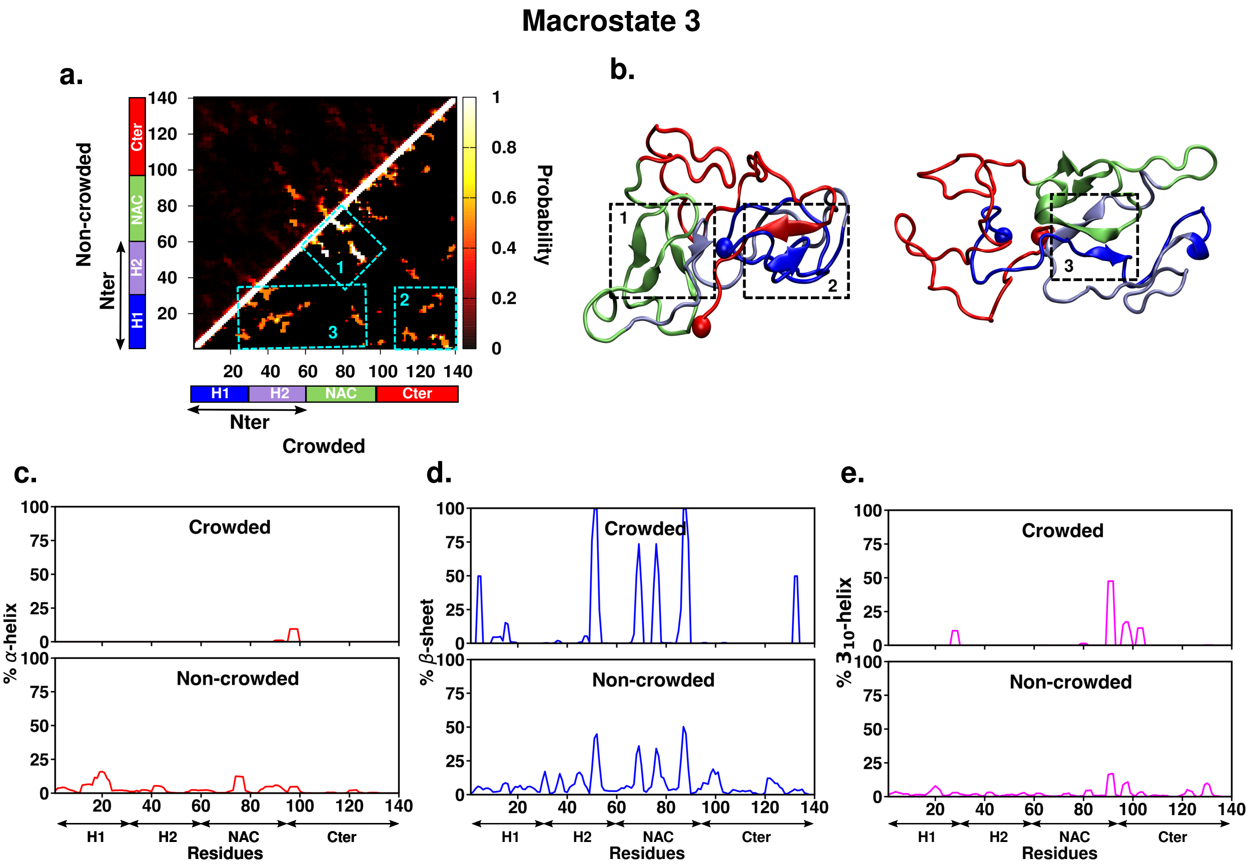


Figure S4: (a)Intrapeptide residue-wise contact probability maps of macrostate MS3 in the absence (upper half of diagonal) and presence (lower half of diagonal) of crowders. The contact probability is indicated by the color scale on the right of the plot (b) Representative conformations from crowding simulation ensemble are shown and colored segment-wise. Dashed boxes in cyan are marked in the crowded system contact map to indicate key contacts and the corresponding secondary structures are marked and numbered in the representative structures. (c) Percentage secondary structure propensity of each residue in the crowded (upper-panel) and non-crowded (lower-panel) systems are presented for (c) $\alpha$-helix (d) $\beta$-sheet and (e) $3_{10}$ helix

Table S1: Simulation details of crowding simulations

| Simulation details | | | | |
|---|---|---|---|---|
| **MSM macrostate** | **Box size (nm)** | **No. of water molecules** | **No. of crowder molecules** | **Total no. of atoms** |
| MS1 Run 1 | 20.89 × 20.89 × 20.89 | 154077 | 3517 | 829353 |
| MS1 Run 2 | 18.75 × 18.75 × 13.25 | 106708 | 2543 | 581437 |
| MS2 Run 1 | 12.64 × 12.64 × 8.94 | 32181 | 780 | 177549 |
| MS2 Run 2 | 10.96 × 10.96 × 7.75 | 20273 | 509 | 113657 |
| MS3 Run 1 | 8.88 × 8.88 × 6.28 | 10213 | 270 | 59077 |
| MS3 Run 2 | 9.37 × 9.37 × 6.62 | 12414 | 316 | 70641 |
